# Competitive exclusion strengthens selection for transmissibility and increases the benefit of recombination for within-host adaptation

**DOI:** 10.1101/2020.06.18.158956

**Authors:** Nathan T. Jacobs, Jeffrey N. Weiser

**Affiliations:** Department of Microbiology, New York University Grossman School of Medicine, New York, NY, USA

## Abstract

Pathogens experience selection at multiple scales, given the need to transmit between hosts and replicate within them. This presents the challenge of cross-scale selective conflict when adaptations to one scale compromise fitness at another, such as mutations that improve transmissibility but make individuals less competitive within hosts. Selection operates differently at these scales, with tight transmission bottlenecks subjecting pathogen populations to genetic drift, and large population sizes within hosts enabling efficient selection for beneficial mutations. Compounding the reduction in diversity by transmission bottlenecks is the occupant-intruder competitive strategy exhibited by some pathogens, where the first variant to colonize a host prevents later arriving variants from contributing to infection, preventing immigration and turning transmission into a “founder takes all” contest. Here, we used multiple modeling approaches to examine how this behavior affects the efficiency of selection for both transmissibility and within- host fitness. We find that in the face of a trade-off, selection for transmissibility is maximized under a tight transmission bottleneck that minimizes within-host competition during colonization. While mutations with increased within-host fitness are favored during within-host replication, an occupant-intruder strategy prevents these mutants from displacing established residents and propagating across the host population, leading to their extinction if they are insufficiently transmissible. Finally, a model of competition on the scale of the host population revealed that competitive exclusion limits the propagation of mutations with improved within-host fitness, unless resident populations can incorporate alleles from intruding variants by recombination. Thus, competitive exclusion may facilitate the improvement and maintenance of pathogen transmissibility, with directional recombination allowing resident populations to mitigate the potential loss of within-host fitness imposed by this occupant-intruder strategy.

**Author Summary:** Transmission is a defining feature of infectious diseases, and so a better understanding of how transmissibility evolves is important for improving disease surveillance and prevention. Successful transmission is often achieved by a small number of individuals which, after establishing residency in a host, may prevent newcomers from participating in infection. Here, we use modeling to examine how competitive exclusion of challengers by resident populations affects the balance between within-host competitive ability and transmissibility. We find that competitive exclusion strengthens selection for transmissibility by disproportionately benefitting the first variant to colonize a host and preventing mutants that may be more competitive but less transmissible from displacing established residents. Competitive exclusion also limits the propagation of mutants that improve within-host fitness without reducing transmissibility, however, increasing the advantage of recombination that allows resident populations to acquire beneficial alleles from challengers. Competitive strategies that allow pathogens to “claim ownership” of hosts may thus help pathogen populations maintain transmissibility, with genetic recombination facilitating within-host adaptation through the incorporation of beneficial alleles from challengers.

## Introduction

Pathogen life history is defined by replication to large population sizes within individuals hosts and the need to transmit to new ones before clearance or host death. Pathogens thus face selection at both the within- and between-host scales, presenting a challenge for evolution when adaptations at one scale reduce fitness at another. Within-host adaptation may be short-sighted, for example, favoring growth in the current host while totally compromising transmissibility [1]. And on the epidemiological scale, selection for transmissibility occurs at the interface between hosts, but mutations that increase transmissibility may be selected against during within-host growth, even if the cost to within-host fitness is mild. This potential for cross-scale selective conflict raises the question of how pathogens can balance both between- and within-host fitness.

Pathogen transmission is often characterized by tight genetic bottlenecks in which substantial diversity is lost between the donor and recipient hosts [2-5]. While contraction in diversity may be the result of strong purifying selection [2, 6], experimental analyses of transmission using neutral markers have shown substantial loss of neutral diversity as well, suggesting that infection is often established by one or a few individual virions or bacterial cells [3, 5]. The loss of genetic diversity at this transmission bottleneck causes both a founder effect, wherein the entire population in a recipient host descends from these few variants, and genetic drift, wherein benefical alleles may be lost and deleterious alleles fixed due to the small size of the founding population.

Both host susceptibility and pathogen behavior can influence the size of the transmission bottleneck and the balance between selection and genetic drift in transmission. If a host is sufficiently resistant to colonization such that most exposures do not result in transmission, then successful colonization could be mediated by rare events in which a single individual establishes infection on its own. This may explain the tight bottlenecks observed in settings with repeated exposures, such as household transmission in humans or between co-housed animals, as the head start achieved by the first variant to transmit puts later arriving individuals at a disadvantage, making it less likely that they will contribute meaningfully to infection in the recipient host [3, 4]. This bottleneck is wider, though, in conditions of increased host susceptibility or closer contact where an exposure comprises more infectious particles, suggesting that transmission is driven largely by the successes and failure of individual pathogens [7, 8].

After establishing infection, a pathogen population may prevent later arriving pathogens from infecting the same host, in a process termed competitive exclusion or competitive exclusion. This phenomenon has been described for multiple taxonomically diverse pathogens, such as influenza A viruses (IAV) and *Streptococcus pneumoniae* [9, 10]. In both cases, co-infection by different genotypes of the same pathogen in animal models can occur only when both arrive at the same time or within several hours of each other. After 18 hours, guinea pigs inoculated with one IAV variant can no longer be superinfected by a second variant, which *in vitro* experiments attribute to the prevention of cellular co-infection [9]. Furthermore, once *Streptococcus pneumoniae* in the mouse nasopharynx reach a quorum after approximately 6 hours of colonization, they begin producing bacteriocins and cognate immunity proteins that later arriving planktonic cells do not, leading to efficient killing of superinfecting bacteria, in a process referred to as fratricide [10]. This turns transmission into a “founder takes all” contest, in which the first variants to arrive become the dominant genotype in the recipient host, and prevents the founder variant(s) from being displaced by a variant that is less transmissible but more fit during within-host competition. This strengthens selection for transmissibility, but also prevents immigration of variants with greater within-host fitness, thereby making within-host adaptation reliant on mutation.

Once infection is established, microorganisms typically grow to large population sizes within the host. In contrast to the genetic drift that may occur during transmission, selection operates more efficiently in these large within-host populations, with within-host adaptation being observed in many human infections with pathogens such as *Staphylococcus aureus* and *S. pneumoniae* [11, 12]. Novel genotypes favored by selection may be generated by mutation during within-host replication, or when the presence of multiple viral or bacterial genotypes within the same host causes genes from one genotype to enter the background of the other. Recombination or reassortment in viruses carries the additional requirement of cellular coinfection, as recombinant genomes are generated by template switching during replication in the cell and reassortant progeny are generated by the packaging of genome segments from different sources. Gene flow between coinfecting bacterial populations, however, may be less restricted, as it can occur through multiple mechanisms such as uptake of free DNA of dead bacteria (transformation), direct exchange between cells through a pilus (conjuation), or hitchhiking of bacterial DNA in bacteriophages (transduction). These processes have been shown to accelerate adaptation in multiple pathogen systems, including Φ6 bacteriophage [13, 14], influenza A virus [15, 16], and *S. pneumoniae* [17, 18]. While competitive exclusion generally prevents superinfecting variants from replicating and contributing to infection directly, the incorporation of alleles from these newcomers by an established resident population may serve as a second method of within-host adaptation.

Here we use multiple modeling approaches to predict how interactions between founder effects at the transmission bottleneck, competitive exclusion, and gene flow affect the efficiency of selection for novel alleles that affect both within-host fitness and transmissibility. We find that competitive exclusion can strengthen selection for transmissibility by imposing tight transmission bottlenecks and preventing displacement by superinfecting variants with greater within-host fitness, and that gene flow enables resident populations to selectively incorporate alleles that improve within-host fitness, thereby balancing between- and within-host fitness.

## Results

We used three models to consider how selection within hosts, during transmission between two hosts, and on the scale of the host population affect the balance of transmissibility and within-host fitness. In the first set of models shown in **Figs 1 and 2**, we use probabilistic models of population genetics to consider how the probability an allele is transmitted or reaches fixation in one host is shaped by its initial frequency, relative transmissibility or within-host fitness, and the size of the transmission bottleneck. In the second model shown in **Fig 3**, we use a branching process model to consider the fate of a novel allele that arises during infection, and how its relative transmissibility and within-host fitness affect its chances of extinction or fixation on the host population scale. In the third set of models shown in **Figs 4 and 5**, we use a compartmental susceptible-infected-susceptible (SIS) model in which multiple variants compete for hosts to consider the balance between transmissibility and within-host fitness on the epidemiological scale. A summary of the parameters, and the models in which they are relevant, is provided in **Table 1**.

**Table 1.**
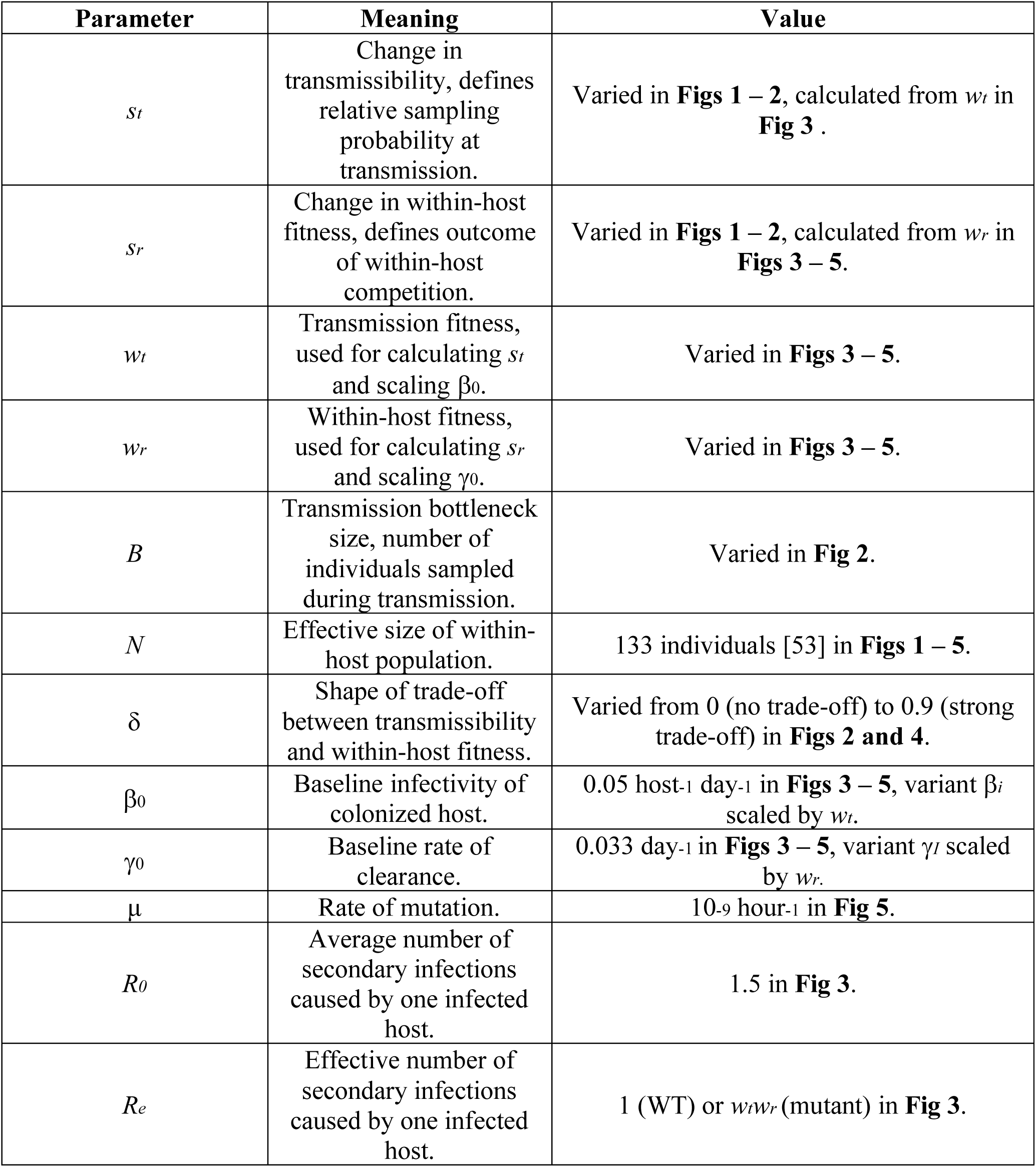
Summary of parameters.

| Parameter | Meaning | Value |
| --- | --- | --- |
| $s_t$ | Change in transmissibility, defines relative sampling probability at transmission. | Varied in **Figs 1 – 2**, calculated from $w_t$ in **Fig 3**. |
| $s_r$ | Change in within-host fitness, defines outcome of within-host competition. | Varied in **Figs 1 – 2**, calculated from $w_r$ in **Figs 3 – 5**. |
| $w_t$ | Transmission fitness, used for calculating $s_t$ and scaling $\beta_0$. | Varied in **Figs 3 – 5**. |
| $w_r$ | Within-host fitness, used for calculating $s_r$ and scaling $\gamma_0$. | Varied in **Figs 3 – 5**. |
| $B$ | Transmission bottleneck size, number of individuals sampled during transmission. | Varied in **Fig 2**. |
| $N$ | Effective size of within-host population. | 133 individuals [53] in **Figs 1 – 5**. |
| $\delta$ | Shape of trade-off between transmissibility and within-host fitness. | Varied from 0 (no trade-off) to 0.9 (strong trade-off) in **Figs 2 and 4**. |
| $\beta_0$ | Baseline infectivity of colonized host. | 0.05 host<sup>-1</sup> day<sup>-1</sup> in **Figs 3 – 5**, variant $\beta_i$ scaled by $w_t$. |
| $\gamma_0$ | Baseline rate of clearance. | 0.033 day<sup>-1</sup> in **Figs 3 – 5**, variant $\gamma_i$ scaled by $w_r$. |
| $\mu$ | Rate of mutation. | 10<sup>-9</sup> hour<sup>-1</sup> in **Fig 5**. |
| $R_0$ | Average number of secondary infections caused by one infected host. | 1.5 in **Fig 3**. |
| $R_e$ | Effective number of secondary infections caused by one infected host. | 1 (WT) or $w_t w_r$ (mutant) in **Fig 3**. |

**Figure 1.**
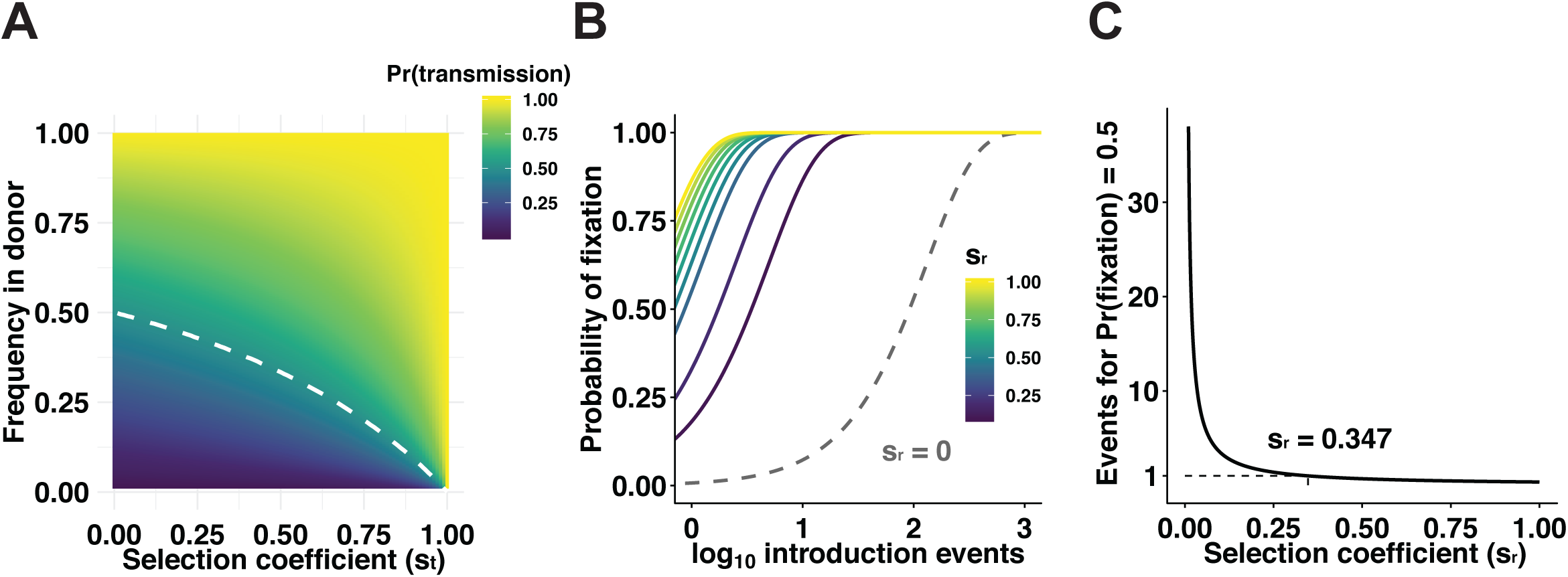
Minority alleles are likely lost during transmission even if beneficial, but alleles that improve within-host fitness are readily selected during infection. The efficiencies of selection for a mutant allele at the transmission bottleneck and following its introduction into a within-host population were considered. A. The probability of transmission was calculated for an allele improving transmissibility with selection coefficient *s*_*t*_ and frequency *p* in the donor host. White dashed line traces the boundary where Pr(transmission) = 0.5. B. The probability of fixation was calculated for a novel allele improving within-host fitness with selection coefficient *s*_*r*_ that is introduced into a within-host population “A times. Gray dashed line represents the probability of fixation of a neutral allele. C. The number of independent introduction events required for an allele with varying *s*_*r*_ to have a 50% chance of fixation was calculated. Dashed line denotes the minimum selection coefficient for an allele to have a 50% chance of fixation after just one introduction.

**Figure 2.**
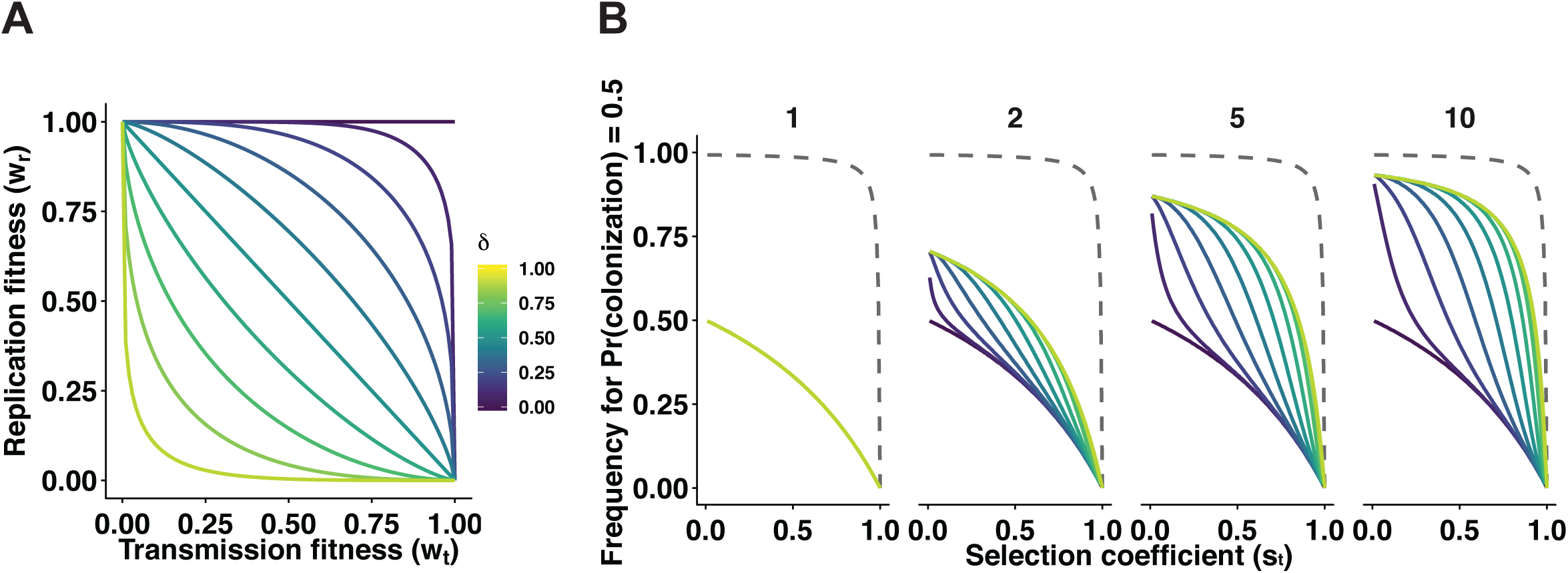
When a trade-off exists between transmissibility and within-host fitness, selection for more transmissible variants is most effective with a tight transmission bottleneck. The effect of transmission bottlenecks on the efficiency of selection for more transmissible variants was considered for various trade-offs between transmissibility and within-host fitness.

A. The shape parameter delta governs the functional form of the trade-off, resulting in a neutral (δ = 0), convex (0 < δ < 0.5), or concave (δ > 0.5) relationship. At higher δ, increases in transmissibility carry greater costs to within-host fitness.
B. The threshold frequency that a mutant with increased transmissibility (expressed by selection coefficient *s*_*t*_) must have in the donor host to have a 50% probability of establishing infection in a recipient host was calculated. For a given transmission bottleneck width (*B*), *B* individuals are sampled from the donor host based on their transmissibility, which then compete for dominance in the recipient host based on their within-host fitness. Gray dashed lines in each facet represent a wide transmission bottleneck of 100 individuals.

**Figure 3.**
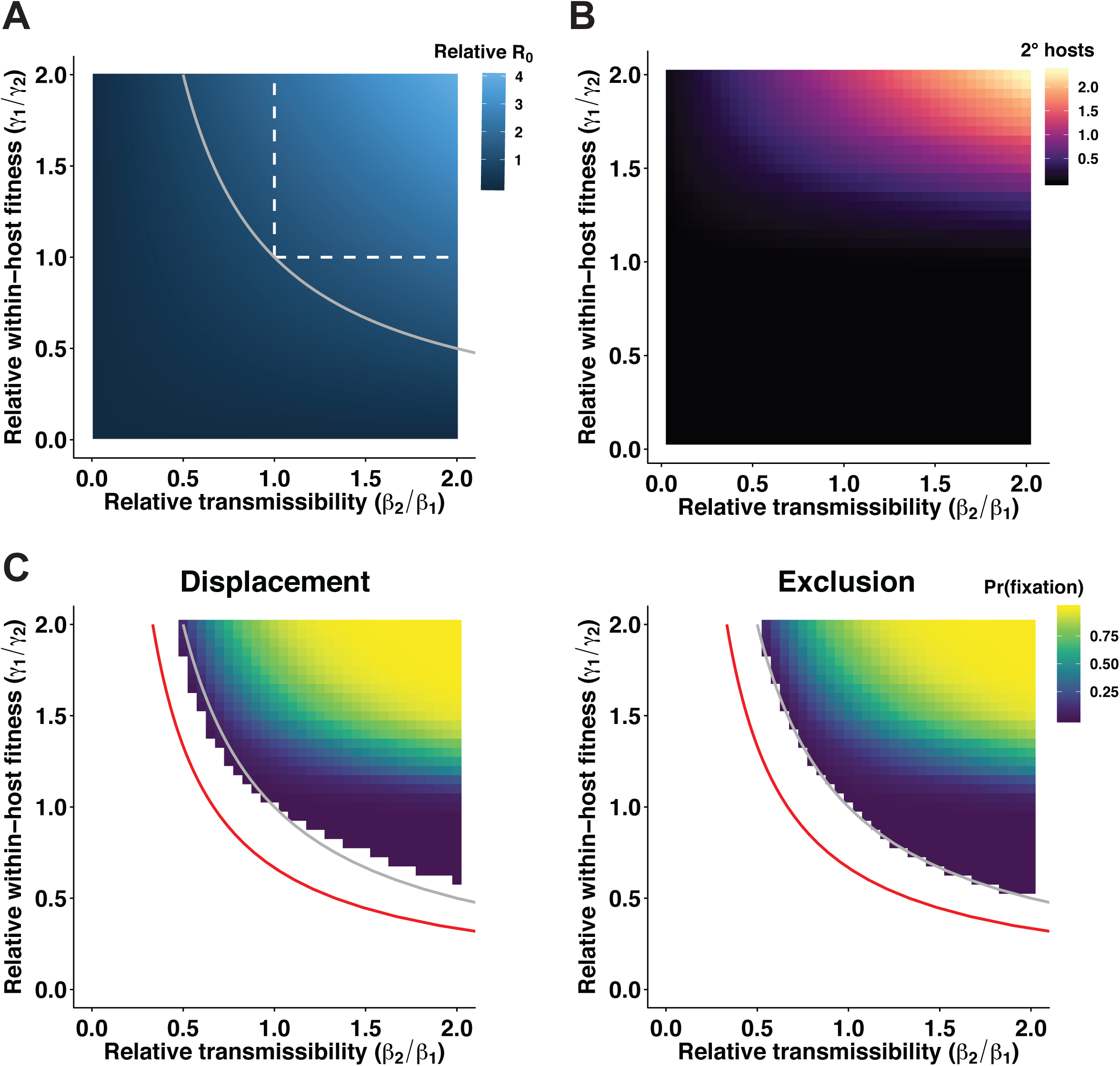
Mutants with increased within-host fitness but reduced transmissibility are likely to be transmitted initially, but competitive exclusion prevents the fixation of mutations that ultimately decrease R_**0**_. For a pathogen population at an endemic equilibrium (R_e_ = 1), the fate of a novel allele with a given effect on both transmissibility and within-host fitness was considered. A. The R_0_ of a mutant, relative to the wild-type ancestor, was calculated based on its relative transmissibility and within-host fitness. Gray solid line represents the boundary below which mutants have reduced R_0_. Regions bordered by the gray solid line and a white dashed line represent mutations that increase R_0_ by increasing within-host fitness at a cost to transmissibility or vice versa. B. The number of secondary mutant infections caused by the host in which a mutation arises was calculated for the mutations shown in a). C. The probability of fixation is shown for mutants that have an R_e_ > 1. As in A), the gray line denotes the boundary below which mutants have a lower R_0_ than wild-type. Red line denotes the boundary below which mutants have R_0_ < 1. In the facet labeled “Displacement,” a variant with greater within-host fitness may superinfect and displace a less fit resident population, while in the facet labeled “Exclusion,” competitive exclusion is perfectly efficient and later arriving challengers cannot contribute to infection.

**Figure 4.**
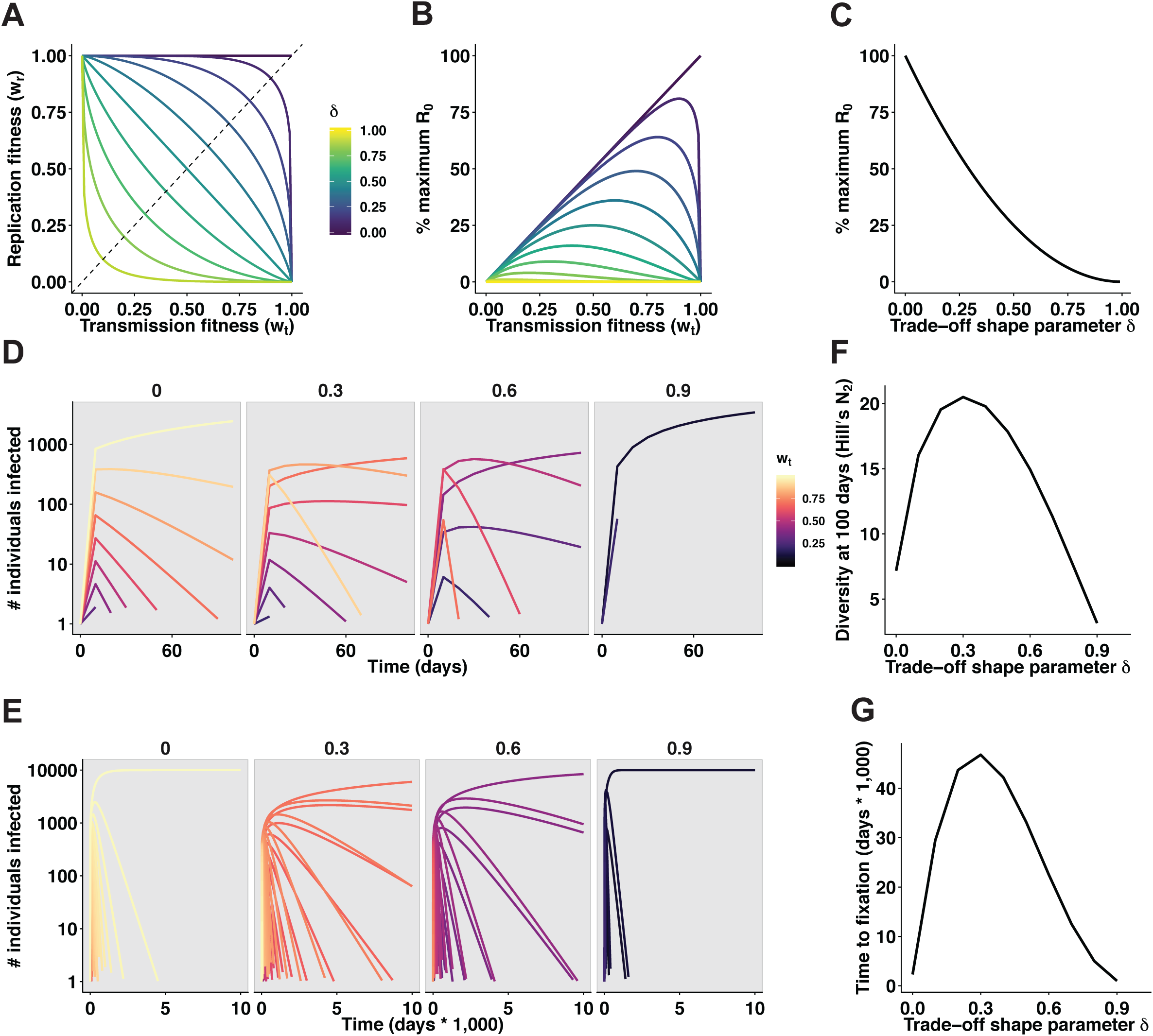
In the face of a transmission-duration trade-off, competitive exclusion allows the early dominance of the most transmissible variants and ultimate fixation of the variant with optimal balance between transmissibility and infection duration. An SIS model was used to analyze the effect of a trade-off between transmissibility and within-host fitness (as measured by infection duration) on competition between variants on the epidemiogoical scale. A. The functional form of the trade-off, as in **2a**, is shown, denoting all possible trait pairs for a given shape parameter δ. Intersection with dashed line (*w*_*r*_ = *w*_*t*_ = 1 – δ) denotes the position of the optimal trait pair for each value of δ. B. The relationship between R_0_, relative to the maximum achievable R_0_ without a trade-off, and transmissibility is shown for a range of δ values. C. The maximum achievable R_0_, relative to the maximum possible without a trade-off, is shown for a range of δ values. D. The prevalences of variants with differing transmissibility are shown during the beginning of an epidemic that begins with 100 hosts, each infected with a different variant. The color of each line represents the transmissibility (lighter color = higher *w*_*t*_) of a given variant, and δ is shown above each facet. For simplicity, only variants with *w*_*t*_ in (0.01, 0.1, 0.2…0.9, 0.99) are shown. E. As in **D**., but all variants are shown for an extended duration. F. The effective diversity, as measured by Hill’s N_2_, of the epidemic at day 100 is shown for a range of δ values. G. The length of time taken for the variant with optimal R_0_ to outcompete other variants and reach 99% frequency is shown for a range of δ values.

**Figure 5.**
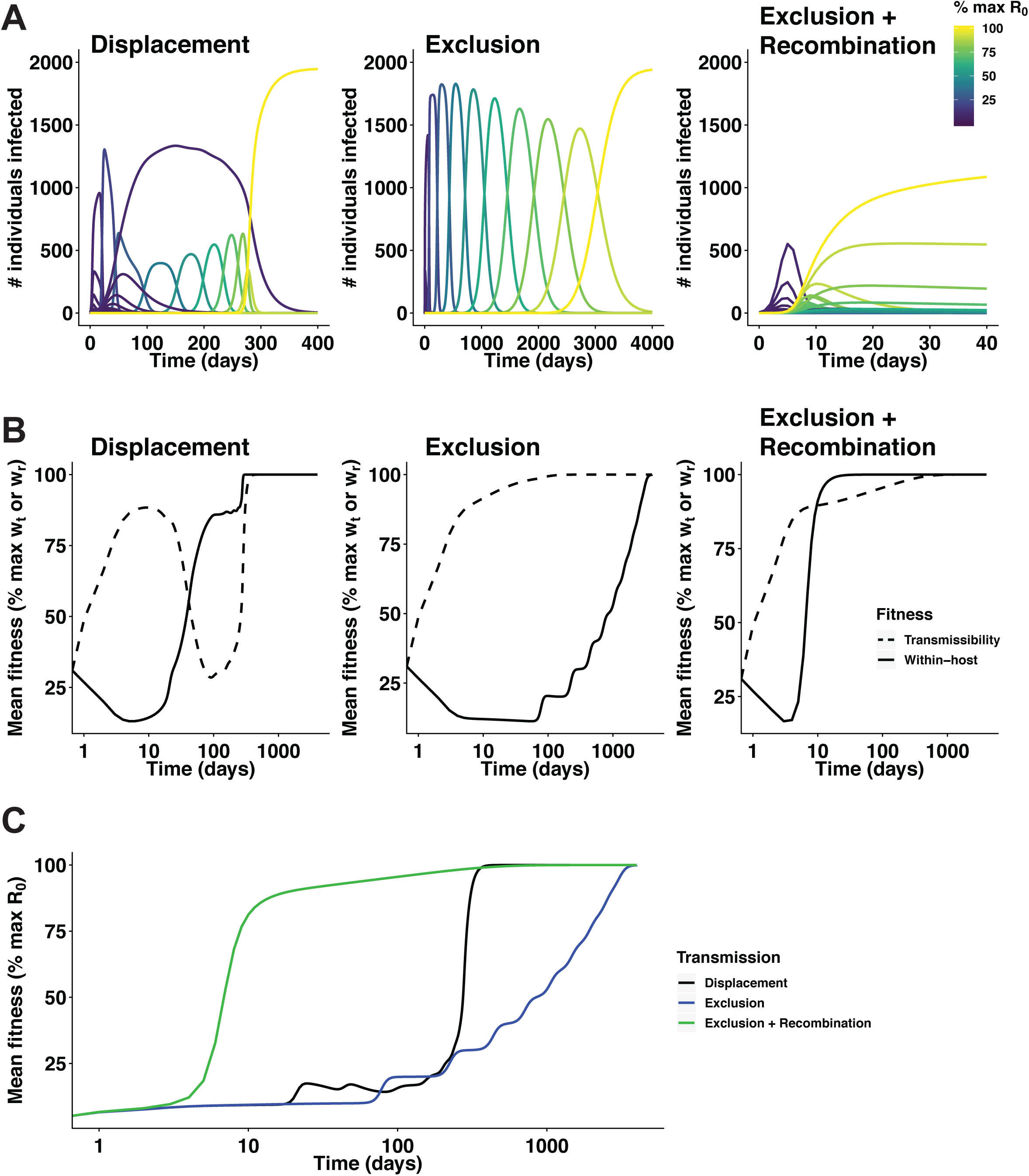
An occupant-intruder strategy for excluding challengers selects for transmissibility, while recombination facilitates the incorporation of alleles that improve within-host fitness. An SIS model was used to examine the evolution of both transmissibility and within-host fitness when no trade-off exists between the two, during an epidemic that begins with some hosts carrying more transmissible variants with low within-host fitness, and some carrying poorly transmissible variants with high within-host fitness. Three different modes of competition govern interactions between two infected hosts: 1) “Displacement,” in which a variant with greater within-host fitness may superinfect and displace a less fit variant through competition, 2) “Exclusion,” in which superinfecting variants are efficiently killed and displacement is impossible, and 3) “Exclusion + Recombination,” in which superinfecting variants are killed, but the resident population may incorporate within-host fitness alleles from challengers by recombination. A. The prevalences of variants is shown over time for three different modes of competition. Lines are colored by the relative R_0_ of a variant compared to the R_0_ of the optimal genotype with maximum transmissibility and within-host fitness. B. The average transmissibility (dashed lines) and within-host fitness (solid lines) of circulating variants are shown over time for each mode of competition. C. The average R_0_ of circulating variants, relative to the maximum achievable R_0_, is shown over time for each mode of competition.

### The efficiency of selection in transmission and the within-host environment

Novel alleles may arise during the course of infection in a host, but must be transmitted to new hosts to avoid extinction. To evaluate the probability of a novel allele being transmitted, we developed a probabilistic model in which the likelihood an allele is transmitted depends on its frequency, *p*, in the donor host, and the extent to which it improves transmissibility, expressed as a selection coefficient, *s*_*t*_. In **Fig 1A**, we examine how both *p* and *s*_*t*_ affect the probability of that an individual variant carrying an allele will colonize the recipient host. When *s*_*t*_ is low, an allele is likely to be transmitted only if it is nearly ubiquitous in the donor population. Even very beneficial alleles with high *s*_*t*_, though, must still be present at moderate frequencies to have an appreciable chance of transmission.

New individuals entering an already colonized host are unlikely to contribute meaningfully to infection, but beneficial alleles from these individuals may be incorporated into a resident population through recombination. To evaluate the conditions under which this process could facilitate the fixation of novel alleles, we considered the probability that a novel allele benefitting within-host fitness with selection coefficient *s*_*r*_ would fix after being introduced to a within-host population a given number of times, whether by recombination or *de novo* mutation (**Fig 1B**). We observe that even slightly beneficial alleles are much more likely to reach fixation upon incorporation than neutral ones, which require ∼100 independent introductions to have a 50% chance of fixing. The required number of introductions required for an allele to have a 50% chance of fixation is shown in **Fig 1C**, revealing that alleles with selection coefficients of at least 0.347 are likely to fix after just one introduction event. These probabilities depend on the effective size of the colonizing population and are therefore sensitive to perturbations by environmental factors such as host immunity or antimicrobial therapies.

### Selection at the transmission bottleneck in the face of a trade-off

Pathogen populations grow to large sizes within a host, but transmission is often characterized by a tight bottleneck in which few individuals colonize a new host. To determine the effect of transmission bottlenecks on the efficiency of selection during transmission, we considered how the size of such a bottleneck affected the probability that an allele would be transmitted, as in **Fig 1A**. In this model, a trade-off exists between transmissibility and within-host fitness, such that increases in transmissibility come at the cost of within-host fitness. The relative fitness of a novel variant is thus described by two distinct selection coefficients, *s*_*t*_ and *s*_*r*_, denoting changes in transmissibility and within-host fitness, respectively. The functional form of this trade-off parallels that considered by Farahpour et al. [19] and depends on a shape parameter δ, which can vary from 0 (no trade-off) to 1 (absolute trade-off between transmissibility and within-host fitness) (**Fig 2A**). As δ increases, increases to transmissibility carry more severe costs, which presents two challenges to selection for transmissibility. First, mutant individuals that are co-transmitted with wild-type ones must then compete for dominance in the new host to establish infection, and so the advantage of a mutant in transmission may be outweighted by a disadvantage in competition during the early stages of colonization. Second, as the mutant allele will be present at low frequency when first generated, the competitive disadvantage faced by the mutant may prevent it from reaching an appreciable frequency in the donor host. We explore this problem in **Fig 2B**, estimating the threshold frequency that an allele with a given selection coefficient *s*_*t*_ must have in the donor host in order to have a 50% probability of establishing colonization in the recipient host for differently sized transmission bottlenecks. For wider bottlenecks, where more individuals compete during the early stage of infection, the advantage of increased transmissibility is dwarfed by the disadvantage of reduced within-host fitness, and transmission mutants require progressively higher frequencies in the donor host to have the same 50% probability of establishing infection in the recipient host. Particularly for high values of δ, the requisite frequency in the donor host approaches the within-host frequency at which all individuals transmitted to the recipient host are transmission mutants, meaning that the allele conferring increased transmissibility is fixed during transmission. Thus, while wide transmission bottlenecks increase the probability that a mutant actually transmits to a recipient host, transmission mutants with reduced within-host fitness benefit the most from tight transmission bottlenecks which allow them to avoid within-host competition in the establishment of new infections.

### Selection in the face of conflicts between scales

In order for a novel allele to emerge in the pathogen population, it must first be transmitted from the host in which it was generated, and from there be propagated across the host population. We model both of these processes to predict the fate of a novel allele based on its transmissibility and within-host fitness relative to those of the wild-type ancestor variant. In this model, “within-host fitness” refers to both competitive ability within a co-infected host (*s*_*r*_ = 1 – *w*_*r*_ for *w*_*r*_ < 1, or 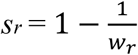for *w*_*r*_ > 1), and the duration of infection when the host is infected with a mutant variant 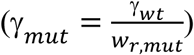. “Transmissibility” refers to both the relative probability that a mutant individual is transmitted from a co-infected host 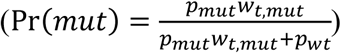 and the relative infectivity of hosts infected with a mutant variant (β*mut* = *w*_*t,mut*_*\**β_*wt*_). The relative R_0_ of this mutant, compared to the wild-type ancestor, is the product of its relative transmissibility and within-host fitness (**Fig 3A**).

The probability that an allele is successfully transmitted depends on both its frequency in that host and its relative transmissibility, as shown in **Fig 1A**. Therefore, we explicity modeled the within-host frequency of a novel allele as a Wright-Fisher process to calculate the average number of new mutant infections caused by the initial host in which the mutation occurred (**Fig 3B**). These simulations show a strong effect of within-host fitness, which helps variants reach higher frequencies shortly after being generated and allows them to cause an appreciable number of secondary infections even if they are less transmissible. Without such an advantage, alleles that are deleterious or even neutral with respect to within-host fitness are less likely to escape the host in which they are generated, as their initially low frequency is not increased by selection. This strong influence of within-host fitness on the initial fate of a novel allele may thus constrain the emergence of mutations that improve transmissibility and ultimately R_0_, especially ones that carry even a slight cost to within-host fitness.

Once a novel variant is successfully transmitted, it may still go extinct if each transmission chain originiating from the initial host ends. To estimate the probability that all transmission from the initial host terminates, indicating extinction, or at least one chain continues indefinitely, indicating fixation of the novel allele in the pathogen population, we used a branching process model (**Fig 3C**). This model calculates the probability that all of an individual’s offspring eventually go extinct based on the expected reproductive success of each individual, which epidemiologically corresponds to the number of secondary cases caused by a single infected individual, R_0_ in a totally susceptible population or R_e_, the effective reproductive number, in a partially susceptible one. When competitive exclusion is perfectly efficient, partitioning wild-type and mutant pathogens into distinct hosts, the R_e_ of a novel variant is simply the product of its relative effects on transmissibility and infection duration, R_e_ = β_*mut*_ * γ _*mut*_. This results in a boundary in which alleles that increase transmissibility at the cost of within-host fitness or vice versa may emerge only if they increase R_0_. If superinfection is allowed to occur, however, than a variant with greater within-host fitness may displace a less competitive resident population from an already colonized host. This increases the relative importance of within-host fitness on the host population scale, reducing the R_e_ of mutants that sacrifice within-host fitness for gains in transmissibility, while increasing the R_e_ of mutants that have made the opposite trade by prioritizing within-host fitness. By extension, this causes some mutants with decreased R_0_ to have R_e_ > 1 and therefore a chance of fixing, while others with increased R_0_ may have R_e_ < 1, preventing fixation. Competitive exclusion thus enhances selection for transmissibility on the host population by 1) limiting the propagation of mutants with reduced R_0_, and 2) preventing more transmissible mutants with increased R_0_ from being displaced by superinfection.

### Competition in the face of a trade-off between transmissibility and infection duration

Traits that promote pathogen transmission adversely can affect within-host survival and vice versa. To determine how a trade-off between transmissibility and within-host fitness shapes epidemiological fitness, we considered a SIS model in which each variant has a distinct transmissibility, *w*_*t*_, and within-host fitness, *w*_*r*_, with the range of possible values being constrained by the trade-off shown in **Fig 2A** (**Fig 4A**). Each of these parameters affects R_0_ in a similar manner to that in the preceding section, i.e., for a given variant *i* with *w*_*t,i*_ and *w*_*r,i*_, β_*i*_ = β_0_ * *w*_*t,i*_ and 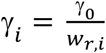, where β_0_ and γ_0_ are the maximum achievable transmissibility and minimum achievable rate of clearance, respectively. The relative R_0_ of a variant is then equal to the product *w*_*t,i*_*w*_*r,i*_, with the maximum R_0_ acheivable for a given trade-off (δ) occurring where *w*_*t*_ = *w*_*r*_ = 1 – δ (**Fig 4B**). As δ increases and the trade-off becomes more concave, the value of this maximum achievable R_0_ decreases (**Fig 4C**).

To determine how these variants that differ in both transmissibility and within-host fitness compete on the epidemiological scale, we analyzed the dynamics of an epidemic in which 100 hosts are initially infected, each with a different variant having *w*_*t*_ from 0.01 to 1 and corresponding *w*_*r*_. We observe that more transmissible variants predominate during the early stages of an epidemic, as they rapidly infect the available susceptible hosts, but are then displaced by the variant with optimal R_0_ (**Fig 4D, 4E**). The dominance of this optimal variant occurs rapidly when δ is close to 0 or 1, but intermediate values, e.g. 0.3 – 0.6, allow prolonged coexistence of variants that are more transmissible or persist for longer, but are ultimately suboptimal, resulting in more diverse epidemics (**Fig 4F**). Similarly, the optimal variant takes longest to reach fixation at intermediate values of δ (**Fig 4G**).

### Balancing between- and within-host fitness by competitive exclusion and recombination

The preceding results demonstrate that competitive exclusion allows a pathogen population to balance transmissibility and within-host fitness in the face of a trade-off between the two, but can also facilitate clonal interference in which beneficial mutations compete with each other. To explore how pathogens might mitigate this detrimental effect of comepetitive exclusion, we considered a SIS model in which variants differ in transmissibility and within-host fitness, but where each aspect of fitness is governed by a separate locus and no trade-off exists between the two traits. Additionally, mutations may occur in infected hosts, altering a variant’s transmissibility or within-host fitness. Finally, interaction between different variants can occur through three different modes of competition. In the first, termed “displacement,” a superinfecting variant with greater within-host fitness may displace a less fit resident variant. In the second, termed “exclusion,” resident variants cannot be displaced by superinfecting variants. In the third, termed “exclusion + recombination,” residents cannot be displaced, but may incorporate alleles from challengers through recombination. The biological mechanism of recombination would depend on the agent in question, such as transformation in bacteria or reassortment in segmented viruses, but it is presumed that alleles from challengers enter the background of the resident population.

To determine how the competitive modes described above affected a pathogen’s exploration of a fitness landscape, we analyzed changes in pathogen transmissibility and within-host fitness over time on the epidemiological scale. The epidemic begins with some hosts infected with variants that have uniformly low within-host fitness but vary in transmissibility, and some hosts infected with poorly transmitting variants that differ in within-host fitness. As in **Fig 4**, we observe that the most transmissible variants quickly predominate as they infect the available susceptible hosts (**Fig 5A**). When displacement by superinfection is allowed, these variants are readily displaced by those with greater within-host fitness, with average transmissibility decreasing as average within-host fitness increases (**Fig 5B**). Emergence of the genotype with maximum fitness in both dimensions is driven by mutations that improve the transmissibility of the most competitive variants. Competitive exclusion allows these more transmissible variants to avoid displacement by more competitive challengers and maintain their early advantage, but emergence of the optimal genotype is conversely driven by mutations that improve within-host fitness. This process is notably slower, as variants with greater within-host fitness can only be propagated by causing new infections. The coupling of competitive exclusion and recombination, however, allows the alleles conferring high within-host fitness to be incorporated onto the background of the most transmissible variants, resulting in a rapid increase in the average R_0_ of circulating variants compared to the other two modes (**Fig 5C**).

## Discussion

We report that cross-scale selective conflict shapes the balance between transmissibility and within-host fitness in two ways. First, the high efficiency of selection for growth within the host allows mutants with greater within-host fitness to predominate, even if they greatly compromise transmissibility. Second, the dependence of a variant’s transmission probability on its within-host frequency hinders selection for mutants that improve transmissibility but not within-host fitness. Left unchecked, this could allow selection to favor variants with greater within-host fitness but lower R_0_, but competitive exclusion prevents the propagation of these mutants on the epidemiological scale. The same occupant-intruder strategy may delay within-host adaptation, however, by subjecting populations to genetic drift in transmission and preventing immigration of variants with greater within-host fitness. Pathogens exhibiting this occupant-intruder strategy can thus adapt to the within-host environment faster by incorporating beneficial alleles from intruders.

A limitation of this study is that our model does not allow for variation in host susceptibility to infection, which may significantly impact the early spread of a mutant. Recent theoretical work by Morris et al., for example, found that infection history creates strong selection at the point of transmission for antibody escape variants [20]. As escape from the first host in which a mutation appears is an important first step in emergence, the probability that such an escape mutant emerges would depend strongly on the local frequency of hosts with those antibodies.

Transmission bottlenecks have previously been studied for many infectious agents such as HIV, IAV, *S. pneumoniae* [2-5, 21]. More general theoretical work predicted that transmission mutants may be favored by tight bottlenecks in organisms, but that such bottlenecks would be rare for directly transmitted pathogens like ones that replicate in the respiratory tract [22]. Our model agrees that in the face of a growth-transmission trade-off, transmission mutants have the greatest chance of establishing new infections under a tight transmission bottleneck that allows them to avoid competition during colonization, but predicts that competitive exclusion may promote selection for transmission even in respiratory pathogens like IAV and *S. pneumoniae*. More recent theory has investigated the influence of transmission bottlenecks on the evolution of an emerging pathogen, specifically with R_0_ < 1, and the probability that a mutation prevents its extinction in the human population [23, 24]. These models, like ours, find that tight transmission bottlenecks can aid selection for transmissibility through a founder effect, and that wide bottlenecks can prevent the emergence of mutations that improve transmissibility at the expense of within-host fitness, even if they improve R_0_. Our model diverges from theirs in that it considers a host-adapted pathogen with R_0_ > 1 at an endemic equilibrium with R_e_ = 1, and so mutants can have a reduced R_0_ that is still greater than one. This allowed us to show that cross-scale selective conflict, in addition to hindering the emergence of more transmissible but less competitive variants as shown by Schreiber et al., can also facilitate the emergence of mutants with greater within-host fitness but ultimately lower R_0_ [24].

In conjunction with previous empirical studies of transmission, our model underscores the importance of transmission in purging variants that result from short-sighted within-host evolution [25, 26]. Deleterious mutations such as defective viral genomes, for example, are consistently generated *in vivo* and replicated with help from abundant cellular coinfection, but transmission is often driven by individual virions, selecting against mutants that are not viable at low MOI conditions [26-29]. Relaxing selection for transmission, however, selects for within-host fitness without consideration for transmissibility. Serial passage experiments in mice, for example, have resulted in the inactivation of genes that promote transmission at the expense of within-host fitness, such as *dltB* in *S. pneumoniae* [30], or selected for accelerated growth in *Plasmodium chabaudi* [31, 32]. Analogous *dlt* inactivation and evolution of more virulent *Plasmodium knowlesi* parasites have also been observed during human infections, suggesting that within-host adaptation has important consequences for disease severity and warranting further study of how transmission affects the rate of such adaptation [11, 33]. Replication by blood-stage *Plasmodium* parasites also involves significant epigenetic changes, which are reversed during passage through the mosquito, further underscoring the importance of transmission in counteracting the potentially detrimental effects of within-host selection [34, 35]. This reversal of short-sighted evolution by transmission does require that transmissible varaints not be completely displaced during infection, suggesting that maintaining diversity in a latent viral reservoir [36, 37] or by continuous genetic exchange during colonization [38, 39] are important for ensuring future transmissibility.

The balance between transmissibility and within-host fitness parallels an invasion-persistence trade-off in other organisms—some genotypes may more readily disperse and colonize new hosts/territories, but are less persistent and thus fare poorly when present in the same environment as less dispersable but more competitive genotypes. This trade-off is evident in several *S. pneumoniae* genes, such as toxin pneumolysin [40, 41], capsule serotype (for example, serotype 4 compared to 23F) [40, 42], and the aforementioned *dltB* [30, 43], which promote transmissibility at the expense of within-host fitness. Whether these genes are favored by selection and maintained depends on the relative importance of transmissibility and within-host fitness, as shown by the loss of *dltB* in serial passage where transmission is more certain. In *Pseudomonas aeruginosa*, selection readily favors a hyperswarming phenotype that doubles the rate of dispersal at a modest 10% reduction in growth rate [44]. By contrast, our results find that selection for comparable transmission mutants with reduced within-host fitness is efficient only once the mutant causes at least one secondary infection, and that the difficulty in escaping the initial host constrains the emergence of such transmission mutants. On the epidemiological scale, the evolutionarily stable genotype tends to be one with intermediate within-host fitness and transmissibility, but the initial abundance of susceptible hosts at the beginning of an epidemic allows the most transmissible variants to enjoy an early advantage. Analyses of spatially structured host populations reveals that these more transmissible strains often dominate the expanding wavefront of an epidemic, similar to the manner in which hyperswarming variants dominate the wavefront of a *Pseudomonas* population [45]. While this early abundance may be transient, it plays an important role in the acquisition of new alleles that benefit within-host fitness without affecting transmissibility—if recombination is directional, with alleles from challengers being incorporated into the resident population, and the resident population is more likely to be a more transmissible genotype, then beneficial alleles from challengers are most likely to enter a more transmissible background.

Our results suggest an important role for directional gene flow in helping resident populations mitigate the effect of genetic drift in transmission and the prevention of immigration imposed by competitive exclusion. It is interesting that the same regulatory circuitry that governs competence for transformation also controls competitive behaviors such as bacteriocin production in *S. pneumoniae* [46] or expression a type VI secretion system in *Vibrio cholerae* [47]. It has been suggested that this shared regulation of competitive behaviors and competence points to a role for neighbor predation in the evolution of these bacteria [48], with recent studies providing empirical support [46, 47]. One argument against the evolutionary benefit of transformation is that the free DNA available for uptake comes from dead cells which, if killed by natural selection, are likely to contribute mutations that reduce fitness [49]. This argument is supported by other studies showing little evidence of recombination driving strongly selected traits such as antibiotic resistance [50], or that DNA acquired through transformation is also useful as a nutrient [51]. However, our results highlight the potential benefits of transformation when a population is not panmictic but structured into distinct hosts—the same occupant-intruder strategy that enchances selection for transmissibility also hinders selection for within-host fitness by imposing genetic drift during transmission and preventing immigration, but recombination allows the incorporation of alleles from challengers onto the background of established resident populations. In contrast to direct competition between a resident and challenger, this directional recombination, in conjunction with efficient selection due to high within-host population size, allows resident populations to selectively retain alleles that improve within-host fitness. Occupant-intruder strategies and recombination may thus play important roles in mitigating cross-scale selective conflicts for infectious agents.

## Methods

### Population genetics of selection at the transmission bottleneck and in within-host populations

The probability that a given allele is transmitted depends on its frequency, *p*, in the population attempting to colonize the susceptible host, and its selection coefficient, *s*_*t*_. The probability that a single individual carrying an allele is transmitted is estimated by:

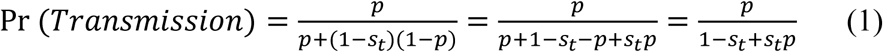

Kimura’s fundamental prediction of allele fixation in haploid populations:

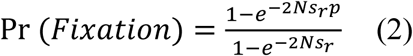

was used to estimate the probability that an allele entering a population by mutation or recombination would eventually reach fixation [52]. *N* was set at a previously estimated effective population size of 133 individuals [53], and ^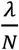^ was substituted for *p* to represent λ individuals acquiring the allele by recombination, yielding the probability of fixation after λ introduction events:

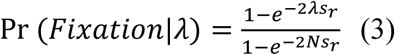

Solving for λ where Pr(fixation) = 0.5 yields the required number of times an allele must be introduced to a population to have a 50% chance of fixation:

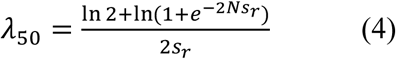

This equation implies that if λ_50 ⩽_ 1, then an allele has a at least a 50% chance of fixation following a single introduction event. Substituting 1 for λ_50_ in (4) yields the inequality

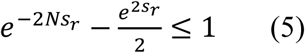

When solved for *s*_*r*_, this inequality reveals the minimum selection coefficient required for fixation to be likely after a single introduction event.

### Fitness trade-offs and population bottlenecks in transmission

When infection is established by multiple individuals, the probability of that an allele is present in the initially colonizing population depends upon the number of individuals transmitted, *B*, and is governed by the binomial distribution:

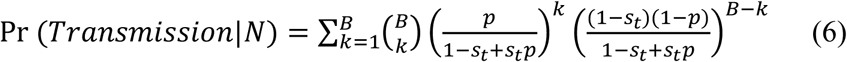

As these initially arriving individuals replicate and compete during the early stages of infection, the probability that an allele will fix in the colonizing population is governed by Kimura’s fundamental prediction of allele fixation in haploid populations [52]:

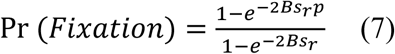

Combining (6) and (7) yields the probability that an individual carrying an allele will establish colonization:

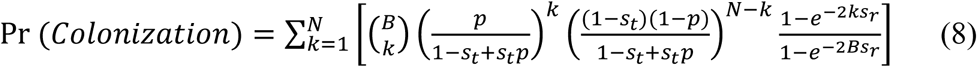

Solving for *p* where Pr(Colonization) = 0.5 yields the required frequency in the donor population for an allele to have a 50% frequency of being transmitted to a susceptible host.

When a trade-off exists between transmissibility and within-host fitness, an allele improving transmissibility with selection coefficient *s*_*t*_ decreases a pathogen’s within-host fitness. The extent of this decrease is governed by the equation:

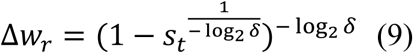

resulting in the negative selection coefficient *s*_*r*_:

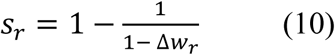

We then refine (8) using these context-specific selection coefficients to estimate the required frequency that an allele must have in the donor host to have a 50% probability of establishing infection in a new host:

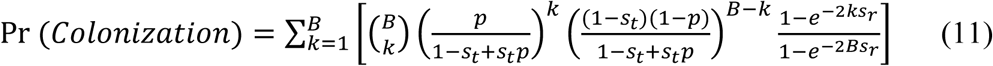

### Branching process model of novel allele emergence

A branching process was used to predict the fate of a novel allele based on its effect on transmissibility and within-host fitness. For a given set of initial conditions, this model calculates the probability that a chain of transmission goes extinct, indicating loss of the novel allele, or continues indefinitely, indicating fixation. In this model, changes in transmissibility affect the infectivity of an infected host 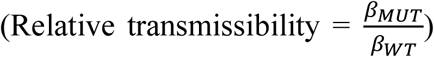, while changes in within-host fitness affect the duration of infection 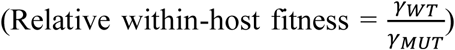, and so the *R*_*0*_ of a mutant relative to the WT ancestor is the product of its relative transmissibility and within-host fitness:

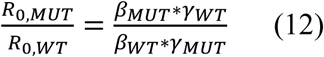

At the within-host scale, changes in transmissibility also affect an allele’s relative probability of being sampled during transmission, as in (1), and changes in within-host fitness affect the within-host frequency of the allele over time. To predict the number of secondary infections caused by a mutant after mutation occurs in the initial host, we simulated the within-host dynamics of infection using a Wright-Fisher process. The mutation arose at a specified time during infection, which varied across a uniform distribution from 12 hours to 29.5 days for an infection lasting 30 days, then fluctuated based on its selection coefficient:

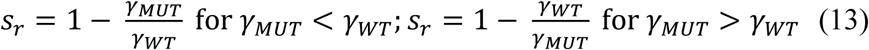

and a generation time of 4 hours [53], for the duration of infection, 30 days. 10 independent simulations per time-of-mutation were conducted. The relative probability of mutant transmission over the course of infection was calculated according to (1), then averaged across all tested mutation times to estimate the number of secondary cases caused by a given mutant after its generation in single host.

This mutation occurs when the number of infected hosts is at an endemic equilibrium and *R*_*e*,_*WT* = 1 due to a reduction in susceptible hosts, so 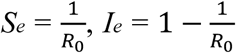, and 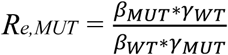. This proportional reduction in susceptible hosts and *R*_*e,MUT*_ assumes that superinfection exclusion is perfectly efficient. To determine how the ability of variants with greater within-host fitness to displace less fit ones through superinfection would affect selection, we modified *R*_*e*,_*MUT* as follows:

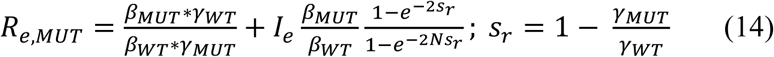

if relative within-host fitness is improved (γ_*MUT*_ < γ_*WT*_), or:

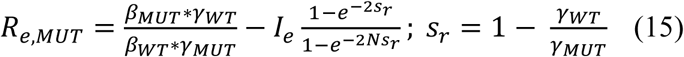

if relative within-host fitness is reduced (γ_*MUT*_ > γ_*WT*_). In both cases, *N* = 133 individuals as before. Based on this reproductive number *R*_*e*,_*MUT* and the expected number of secondary infections caused by the initial host, the extinction probability of a given chain of mutant transmission was estimated using a Galton-Watson process [54]. After the probability a given transmission chain’s extinction was calculated, the probability of ultimate extinction, in which all transmission chains arising from the initial host, was calculated as:

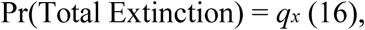

where *q* is the probability of extinction of a transmission chain arising from a single host, and *x* is the number of mutant infections caused by the initial host where the mutant arises. Finally, the probability that at least one transmission chain continued indefinitely, indicating fixation of the mutant in the population, was given by:

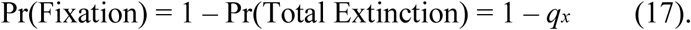

this model is sensitive to the R_0_ of the wild-type ancestor, which here was set at 1.5.

### SIS model with invasion-persistence trade-off

To investigate how trade-offs between transmissibility and within-host fitness affect the selection for both traits on the scale of the host population, we define a Susceptible-Infected-Susceptible (SIS) model using a system of ordinary differential equations (ODEs):

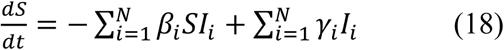

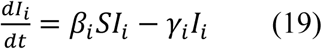

*I*_*i*_ indicates the number of individuals infected with pathogen variant *i*, which has a transmission fitness *w*_*t*_ and corresponding *w*_*r*_ as calculated by the following:

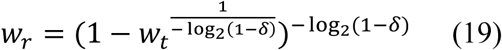

A pathogen’s w_t_ and w_r_ affect its transmissibility and the duration of infection, respectively:

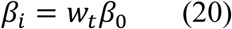

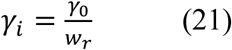

where β_0_ is the maximum achievable transmission coefficient, and γ_0_ is the minimum achievable rate of clearance. The transmission fitness of each type, *w*_*t*_, ranges from 0.01 to 0.99 in increments of 0.01, for a total of 99 types. The system was initialized with 1 infected individual of each type, and 9,901 susceptible individuals, for a total population size of 10,000 hosts. The dynamics were then realized using the R package *deSolve*.

### SIS model with within-host competition, competitive exclusion, and recombination

To determine how superinfection exclusion and recombination affect selection for both transmissibility and within-host fitness, we define an SIS model in which the two traits are governed by independent loci, with *I*_*tr*_ denoting an individual infected with a pathogen of *w*_*t*_ = *t* and *w*_*r*_ = *r*. The change in susceptible individuals is the same as (18):

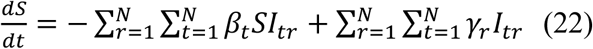

The change in each class of infected individuals follows one of three ecological regimes, referred to herein as “displacement,” “exclusion,” and “exclusion + recombination.” Where resident pathogens may be displaced by superinfecting pathogens that are more fit (“Displacement”):

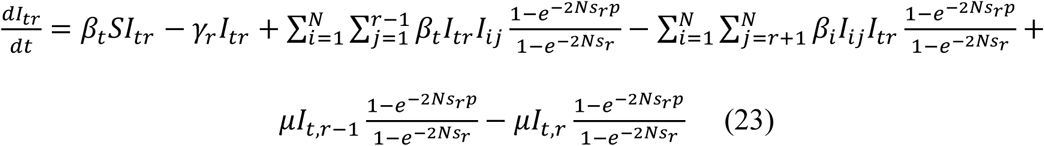

In the case where superinfection exclusion occurs, and resident pathogens exclude superinfecting individuals:

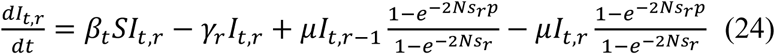

Finally, in the case where superinfection exclusion occurs, but DNA from challengers is available for uptake by the resident population:

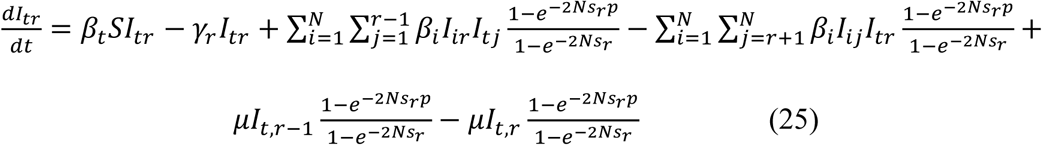

Possible *w*_*t*_ and *w*_*r*_ values ranged from 0.1 to 1 in increments of 0.1, as well as 0.01. The system was initialized with 2,000 hosts: 1,895 susceptible hosts, 5 hosts infected with pathogens of each *w*_*t*_ type and *w*_*r*_ = 0.1, and 5 hosts infected with pathogens of each *w*_*r*_ type with *w*_*t*_ = 0.1, so that the alleles conferring efficient transmission and replication are present initially, but in different hosts. The dynamics were then realized using the R package *deSolve*.

## Acknowledgements

The authors thank John Lees for helpful discussion and review of the manuscript.

